# Ecological feedback on diffusion dynamics

**DOI:** 10.1101/254136

**Authors:** Hye Jin Park, Chaitanya S. Gokhale

**Affiliations:** Department of Evolutionary Theory, Max Planck Institute for Evolutionary Biology, August Thienemann Str. 2, 24306, Plön, Germany; Research Group for Theoretical Models of Eco-evolutionary Dynamics, Department of Evolutionary Theory, Max Planck Institute for Evolutionary Biology, August Thienemann Str. 2, 24306, Plön, Germany

**Keywords:** eco-evolutionary dynamics, social dilemma, spatial dynamics, pattern formation

## Abstract

Spatial patterns are ubiquitous across different scales of organization. Animal coat pattern, spatial organization of insect colonies, and vegetation in arid areas are prominent examples from such diverse ecologies. Typically, pattern formation has been described by reaction-diffusion equations, which considers individuals dispersing between sub-populations of a global pool. This framework applied to public goods game nicely showed the endurance of populations via diffusion and generation of spatial patterns. However, how the spatial characteristics, such as diffusion, are related to the eco-evolutionary process as well as the nature of the feedback from evolution to ecology and vice versa, has been so far neglected. We present a thorough analysis of the ecologically driven evolutionary dynamics in a spatially extended version of ecological public goods games. We show how these evolutionary dynamics feedback into shaping the ecology thus together determining the fate of the system.

## 1 Introduction

Evolutionary game dynamics has been successfully used to describe the evolution of types in a population, be it frequencies of alleles in a biological setting or languages in a cultural setting [1, 2]. The most widely used application of this theory is in addressing social dilemmas. Social dilemmas occur when the interests of an individual and that of the group it belongs to, collide [3]. The ubiquity of social dilemmas is evident by its appearance in pertinent issues such as fishery and wildlife management [4] and global climate change [5]. Biologically relevant scenarios such as foraging strategies [6], group hunting behavior [7, 8], and bacterial secretions interpreted as public goods [9] provide these dilemmas a sociobiological setting. The resolution of social dilemmas lies at the heart of achieving a transition in the level of organization e.g. evolving multicellularity [10] (or the deconstruction of sociality, as in cancer evolution [11]). A number of ways of resolving such dilemmas, elegantly captured by public goods games (PGG), have been proposed [12, 13]. One of the means of resolving public goods games is the imposition of spatial structure on the evolving population. Conceptually, classical ideas such as Wrights island model [14], the haystack model [15], contemporary group selection models [16], evolutionary dynamics with structure and many more [17, 18, 19, 20], impose a condition limiting encounters between the interacting agents. Spatial dynamics thus has been successful in resolving the social dilemma with the possibility to maintain a mixture of cooperators and defectors in the long run [21].

Besides stabilising cooperation, spatial dynamics also results in brilliant spatial patterns under eco-evolutionary processes [22]. Ecological dynamics are incorporated by explicitly accounting for the feedback of population densities on the evolutionary processes and vice versa. We deviate from the classical use of diffusion as a “constant” and investigate an eco-evolutionary feedback on population mobility. Through this novel outlook, it is possible to explain how populations can avoid extinction in a spatially extended selection-diffusion system. This improves our understanding of the ecological aspects of the diffusion process responsible for the spatial patterns. In this manuscript we focus on density dependent diffusion as an ecological factor of spatial dynamics and examine its effect on pattern formation. Especially, we focus on two distinct density dependent dynamics: in growing bacterial cultures and human migration. We establish a connection between the exact properties of diffusion and the observed spatial patterns. The details of the diffusion rule, which can differ between species is shown to be crucial in determining the observed patterns.

## 2 Model and Results

### 2.1 Eco-evolutionary dynamics with diffusion

In PGG, cooperators invest a fixed amount *c* into a common pool. For *m* such cooperators, this common pool of value *mc* is then multiplied by a factor *r* > 1. The benefits of this interaction are returned equally to all individuals *N*, thus *rmc*/*N*. While this is the payoff of a defector, *P*_*D*_(*m*) = *rmc*/*N*, a cooperator, having paid the cost, gets *P*_*C*_ (*m*) = *P*_*D*_ (*m*) – *c*.

In order to incorporate eco-evolutionary dynamics, (normalized) densities are introduced instead of frequencies of cooperators and defectors. Within a patch, the sum of cooperator and defector densities, *u* and *v*, lies between zero and unity, 0 ≤ *u* + *v* ≤ 1, indicating the total density of the population in the patch. Hence, the available space in the patch is denoted by *w* ≡ 1 − *u* − *v*. The group size within each patch is fixed to *N*, but the densities change in time thus affecting the effective group size. Without diffusion between subpopulations, the population dynamics within each patch can be analyzed separately as an independent population [23].

Individuals have a chance to meet another individual with a probability that is proportional to the total density in a well-mixed population. If the population density is small, individuals meet less often and hence form smaller groups. If the density is high then the maximum group size *N* can be reached. As a consequence, the game-interaction group size *S* depends on the total density. The expected payoffs for defectors and cooperators are calculated from the densities of cooperators and defectors. Using the expected payoffs and the probability *p*(*S*; *N*) of observing a group of size *S*, given a maximum group size *N*, the average payoffs for defectors and cooperators, *f*_*D*_ and *f*_*C*_, can be expressed as follows,

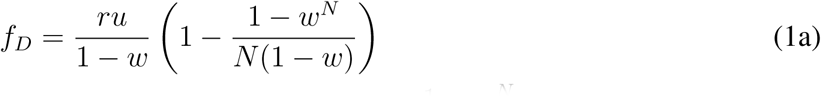

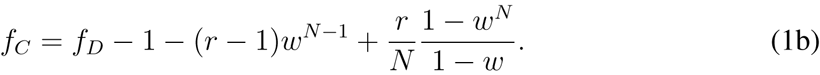

We have set the investment cost *c* = 1 (for details see Appendix.).

If there is available space (*w* > 0), individuals reproduce according to their average payoffs. All individuals are assumed to have the same constant birth and death rates given by *b* and *d*, respectively. The change in the densities of cooperators and defectors over time is captured by the following extension of the replicator dynamics [23, 24, 25],

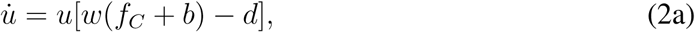

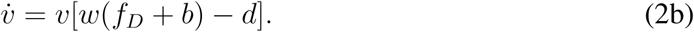

This set of deterministic equations provides us with the time evolution and the fixed points of the system. The asymptotic behavior of the system is determined by the stabilities of these fixed points. The system undergoes a Hopf bifurcation as *r* varies. For small *r*, extinction (*u* = *v* = 0) is the stable fixed point while for a large *r* coexistence (*u*, *v* > 0) becomes stable.

Hence, both cooperators as well as defectors die out for a small rate of return from the public good (*r* < *r*_*hopf*_). With spatial dynamics, however, the stability of the fixed point can change. By forming patterns, cooperators and defectors can coexist even for *r* < *r*_*hopf*_. With diffusion coefficients *D_c_* and *D_d_* for cooperators and defectors, respectively, Eqs. [2] can be rewritten as follows:

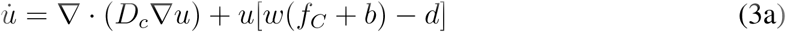

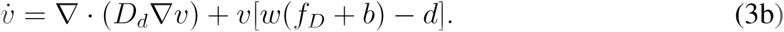

There is no in‐ or out-flux at the boundaries. To avoid confounding effects, there are no game interaction between patches, but only exchange of individuals. According to the constant ratio of diffusion coefficient *D* = *D_d_*/*D_c_* > 1, various patterns have been observed in [22] (see Fig. 1). In the phase space of *r* and *D*, various phases are observed ranging from homogeneous coexistence to extinction via pattern formation. Between the extinction phase and diffusion induced coexistence, chaotic patterns are observed.

**Figure 1:**
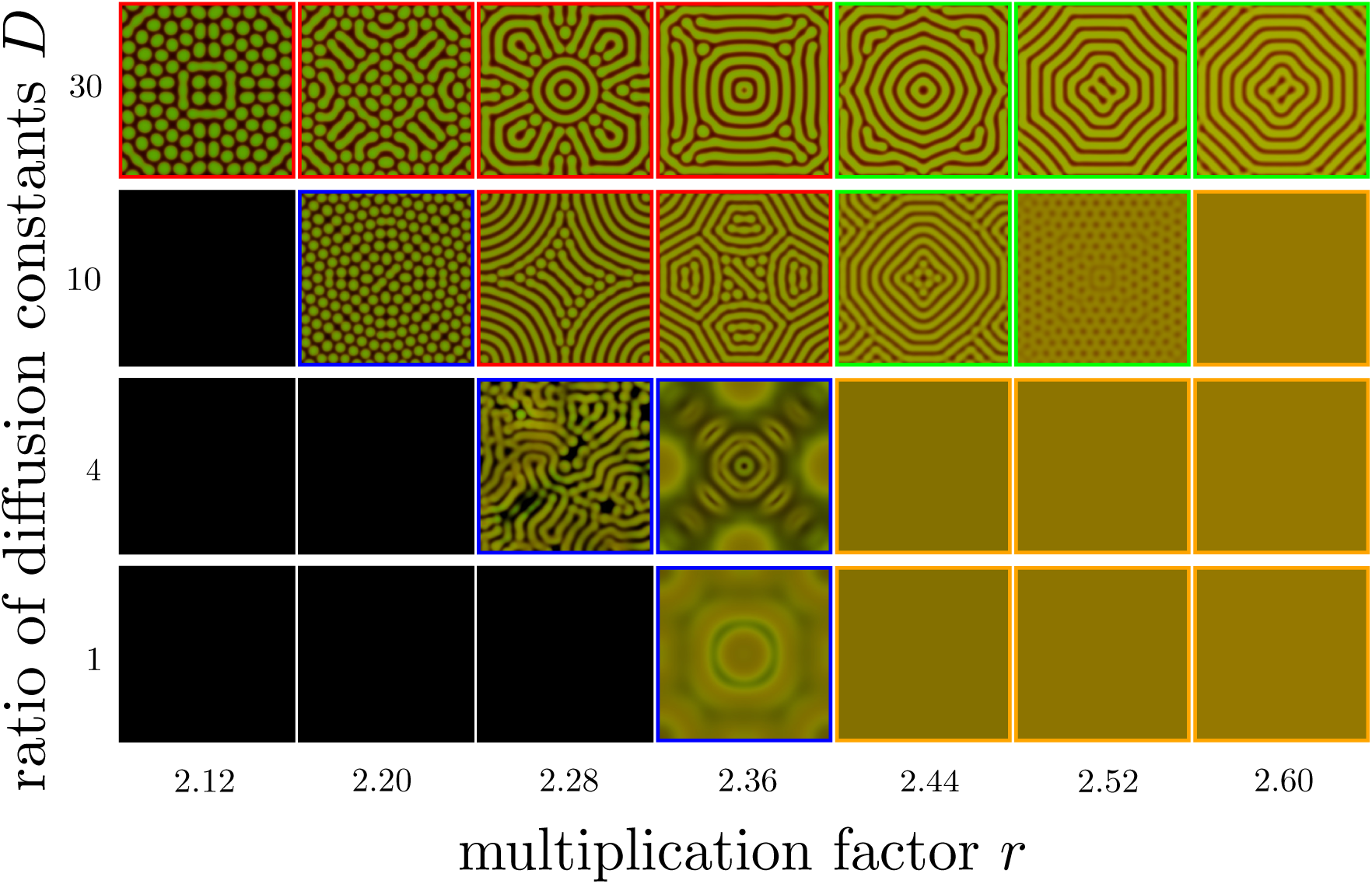
Spatial patterns with various parameters [Reproduction of the figure in [22]]. A two-dimensional square lattice is considered. Each subpopulation resides in a patch where they play the eco-evolutionary public goods game with a maximum group size of *N* = 8. Constant birth rate of *b* = 1 and death rate of *d* = 1.2 are used. Cooperator and defector densities are presented as green and red colors and the brightness indicates the total density (see Appendix). There are five phases (framed using colors), extinction (black), chaos (blue), diffusion induced coexistence (red), diffusion induced instability (green), and homogeneous coexistence (orange). Both diffusion induced coexistence and instability show striped patterns but different regime: first one is for *r* < *r_hopf_*, and the second one is for *r* > *r_hopf_*. The Hopf-bifurcation point is at *r_hopf_* ≃ 2.3658. We used the Crank-Nicolson method to get patterns with a linear system of size *L* = 283. An uniform disk with densities *u* = *v* = 0.1 at a centre is used for an initial condition. Note that the symmetry breaking for *r* = 2.28 and *D* = 4 arises from numerical underflow [26].

### 2.2 Ecological feedback on diffusion dynamics

Density dependent diffusion is observed across scales of organization from microbial systems to human societies [27, 28, 29, 30, 31]. Furthermore, within a population, different type of individuals can show different mobility for example as in the aphid and planthopper populations [32, 33, 34, 35, 36, 37]. We explore the possibility that cooperator and defector densities affect the diffusive behavior of the population. For the sake of simplicity, we assume a fixed diffusion coefficient for cooperators, and develop the defector’s density dependent diffusion. Various instances demonstrate how defectors moving faster than cooperators, survive harsh environment [38, 39], and it will be more interesting to focus on *D* > 1. We investigate the effects of two different dynamic diffusion functions on spatial patterns, one representing the way bacteria diffuse on a petri dish and the other typifying human migration patterns [30, 40].

For the density dependent diffusion of bacteria, we start from a bacterial growth experiment and its modeling [29, 40]. In this model, bacteria grow by consuming nutrients, and spread by diffusing in space. From the experimental result, we interpret that bacteria move faster when they are more productive and slowly when nutrients or bacterial concentrations are low. We thus write the defectors mobility inferred from the productivity of defectors. In Eq. [A.7], the productivity of defectors is determined by *vw*(*f_D_* + *b*). Hence we design the defector’s diffusion coefficient as follows:

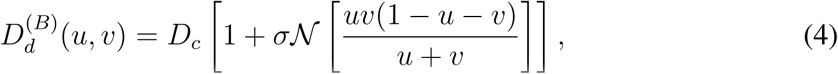

where

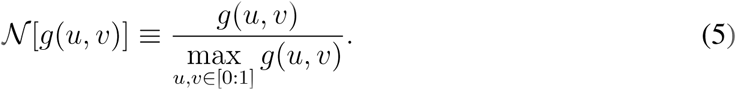

By using the expression 
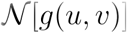
 we put bounds on the range of density dependence from

zero to unity. Here, we use a baseline diffusion value of *D_c_* and focus on *D* > 1. This is done by setting up the diffusion coefficient such that for *u* = *v* = 0 we have 
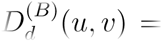
 *D_c_*. The parameter *σ* is the *intensity of density dependence*. As shown in the Fig. 2 (a), the diffusion coefficient 
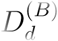
 has a maximum at intermediate densities of *u* and *v*. If the density of cooperators or defectors is too high or too low then they move slowly. At a fixed defector density *v*(*u*) diffusion coefficient is concave in *u*(*v*).

**Figure 2:**
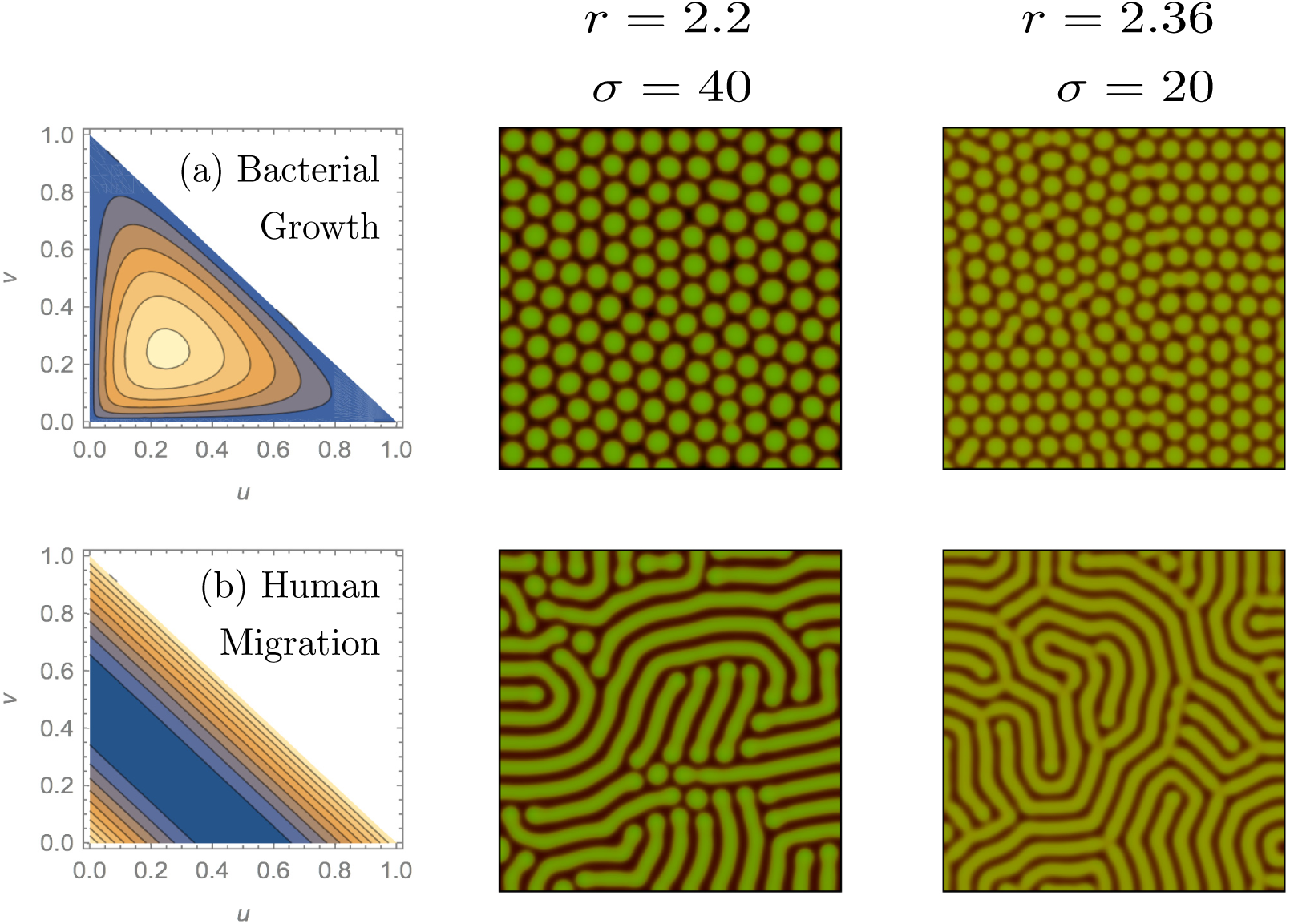
Patterns under the two density-dependent behaviors for *r* = 2.2 and *r* = 2.36 (in column) with *σ* = 40 and *σ* = 20, respectively. The sensitivity to density dependence *σ* are denoted with the multiplication factors *r*. The patterns in the same row as (a) are the result of the density dependent diffusion function as in [4]. The second row (b) is the results of using [6]. The functions *D_d_*(*u*, *v*) are shown as contour plots in *u* and *v* space. Blue and yellow colors represent low and high values at a given *u* and *v*, respectively. For bacterial growth, dotted patterns emerge, while striped patterns appear for human migration. The results show that the fast movement of the defectors in high reproduction region forces the system to the edge of extinction. We use *L* = 283, *N* = 8, *dx* = 1.4, *b* = 1, and *d* = 1.2. Cooperator and defector densities in each patch are randomly drawn from 0 to 0.1 as an initial condition.

In contrast to bacteria, human mobility is maximized at low and high population densities. Utility in humans seems to be maximised by avoiding extremely low and extremely high total population densities [30]. We introduce this diffusion dynamics for defectors by constructing a coefficient as,

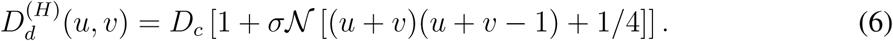

Similar to the bacterial diffusion coefficient, we can divide the human migration coefficient into two parts, density independent and dependent. Here, *σ* again indicates the intensity of density dependence. To keep *D_d_* > 0, we add 1/4 to the second term in Eq. [6]. For the numerical simulations, Crank-Nicolson method is used for constant diffusion coefficient, however we use the forward Euler method for a dynamic diffusion coefficient. Since we are varying *σ*, we focus on *r*-*σ* space instead of *r*-*D* space as opposed to [22]. The diffusion coefficient is not homogeneous in space anymore as it varies from patch to patch. An average diffusion coefficient 
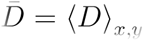
 is determined for a given *σ* and the associated density dependent dynamics.

Diffusion dynamics as described above, is a function of both, evolutionary (fraction of cooperators) as well as ecological (total population density) parameters. This eco-evolutionary diffusion dependence forms the nucleus of our model elucidating the effects of eco-evolutionary processes on pattern formation.

## 3 Analysis and Discussio

We examine patterns for various *r* and *σ* under a varying diffusion coefficient. The different diffusion dynamics for different organisms results in different spatial patterns. As shown in the Fig. 2, bacterial growth gives the dotted patterns which are close to the extinction phase while human migration shows striped patterns which are far from the extinction. Cooperators can survive and resist the invasion of defectors by forming alliances. In spatial terms this translates into minimising the surface area of the colony. Since high cooperator fraction is essential for populations to persist, dotted patterns (with high cooperators fraction) are observed near the extinction phase. Therefore, the emergence of dotted patterns for high multiplication factor and intensity implies that the density-dependent diffusion drives the system to the margins of harsh environment for surviving. This shift towards the margins comes with the risk that along this edge, sometimes, the dynamics can descend into chaos. Surely, this can result from the diffusion property itself. The defectors grow, reproduce and spread their offspring fast, and it induces a fast decrease of cooperators while increasing the risk of extinction. On the other hand, human migration case produces striped patterns which implies that diffusion increases the size of the surviving region in the parameter space.

To confirm our results, we explore the patterns in the parameter space with two functions defined for the defectors diffusion dynamics. As shown in Fig 3, bacterial diffusion mainly forms dotted pattern for *r* < *r_hopf_*, while human migration shows striped patterns. The tendency towards these patterns remains stable even when average diffusion coefficients 
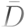
 are the same for both cases. These patterns support that two different density dependent diffusions modify the ranges of the surviving area in the *r*-*σ* parameter space. At a given *σ*, human migration dynamics is more resilient against extinction than bacterial diffusion. Note that this tendency becomes opposite for *r* > *r_hopf_* because two different density dependent diffusions show opposite behavior when *r* increases (see Fig. 4). For small values of *r*, bacterial growth diffusion suppresses average diffusion coefficient 
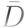
 while human migration diffusion boosts 
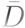
 However, it is opposite for large *r* as shown in Fig. 4. As a result, bacterial diffusion reduces the size of the surviving region for *r* < *r_hopf_* while for *r* > *r_hopf_* diffusion increases the size of the region with patterns as compared with human migration dynamics.

**Figure 3:**
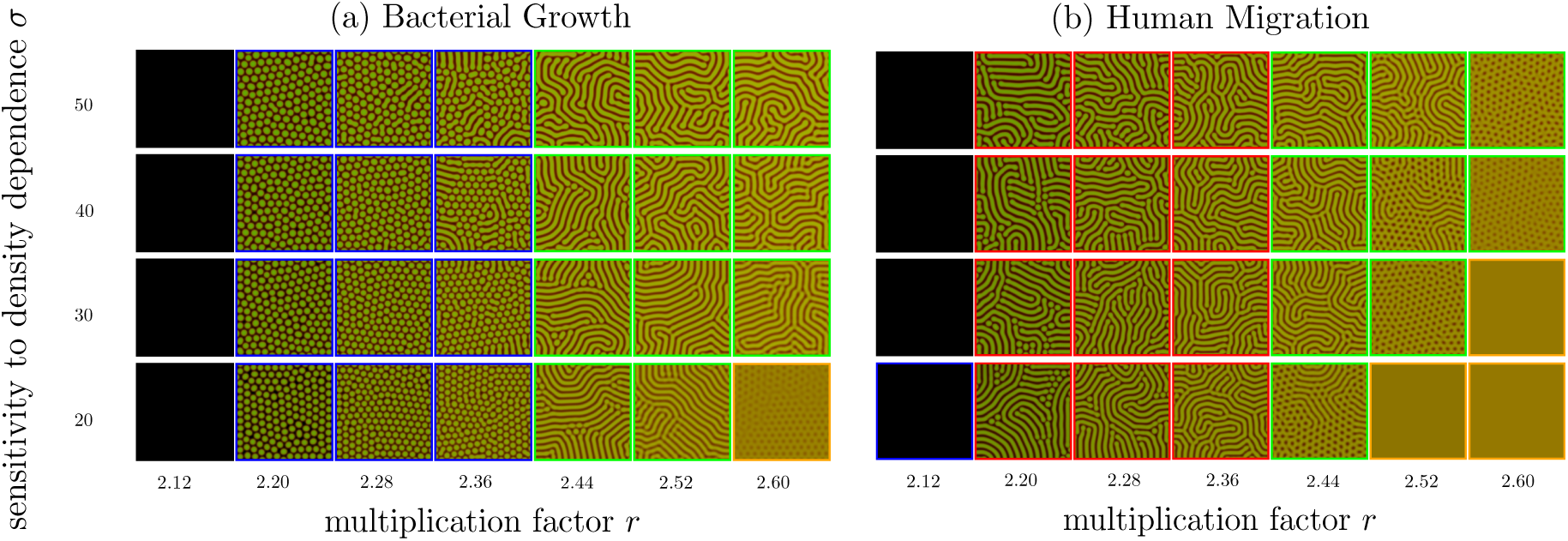
Phase diagrams with different diffusion functions: (a) where the diffusion reflects bacterial growth (4) and (b) where the diffusion reflects human migration (6). Each frame of pattern is colored by the same criteria with Fig. 1 except dotted pattern for *r* < *r_hopf_*. To distinguish dotted and striped patterns for *r* < *r_hopf_*, we have used different colors. For dotted patterns, we used the blue color which is used for chaotic patterns, as both are observed near extinction phase. We can clearly see the different patterns for different density dependent diffusion for *r* < *r_hopf_*. Cooperator and defector densities in each patch are randomly drawn from 0 to 0.1 as an initial condition.

**Figure 4:**
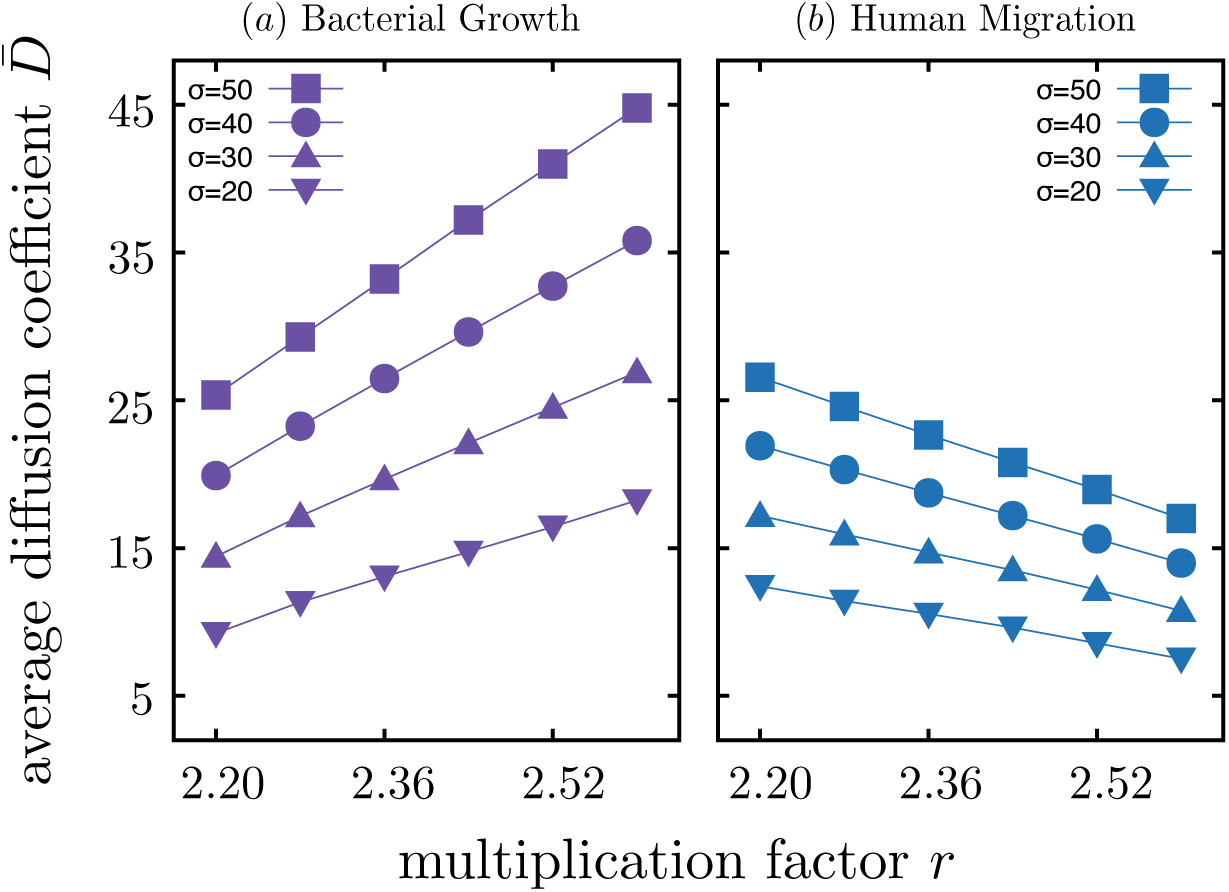
The average diffusion coefficient 
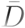
 against the multiplication factor *r* at a given *σ*. Different point symbols represent results of 
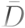
 with different *σ*. Configurations in Fig. 3 are used for calculating 
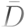
 Solid lines are guide lines for points with the same *σ*. The average diffusion coefficient 
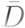
 increases as *r* increases for bacterial growth diffusion, while 
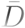
 decreases for human migration diffusion. This opposite behavior over *r* of two different diffusions makes different shifts of two boundaries between heterogeneous and homogeneous pattern phases for *r* < r_hopf_ and r > r_hopf_. For *r* < *r_hopf_*, the boundary under bacterial growth diffusion is located much upper than human migration diffusion, while it is opposite for *r* > *r_hopf_*.

So what property of the density dependent diffusion exactly induces dotted or striped patterns for *r* < *r_hopf_* ? To understand the intricacy of the functions we examine several diffusion coefficient formulas based on the geometries of their key variables. From the Eq. [4] we get an intuition for these important variables. If we transform cooperator and defector densities *u* and *v* to cooperation fraction *f* and total density *ρ*[23, 41], we can further simplify the diffusion coefficient *D_d_* as a function of 
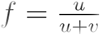
 and *ρ* = *u* + *v*

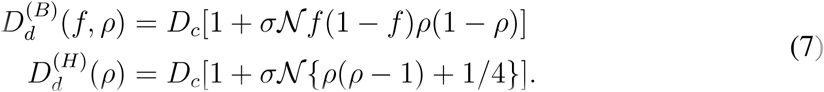

It implies that the key variables can be *f* and *ρ*. These two quantities are exactly the evolutionary and the ecological parameters which are of interest in eco-evolutionary game dynamics [23]. The diffusion coefficients are not just density dependent but can be interpreted as eco-evolutionary diffusion coefficients. To study the impact of these two quantities, we examine all possible combination of *f*, 1 − *f*, *ρ*, and 1 − *ρ* taking into account their geometry. In all, 16 cases have different geometries, and thus they cannot span each others. Density dependent functional forms are drawn in the Fig. 5 (a) in *f* and *ρ* space. Each function is then a product of the respective row and column of Fig. 5 (a). The top left corner is the bacterial diffusion behavior. For patterns obtained for *r* = 2.32 < *r_hopf_*, we compare the results in Fig 5 (b). Without *ρ* in *D_d_*(*f*, *ρ*), striped patterns appear. We can conclude that moving slow in low density *ρ* induces the dotted patterns. However in presence of 1 − *f* we recover striped patterns.If we compare patterns from (1 − *f*)*ρ*(1 − *ρ*) and *ρ*(1 − *ρ*), we can find that fast movement in low cooperator fraction and slow movement in high cooperator fraction recovers striped patterns.

**Figure 5:**
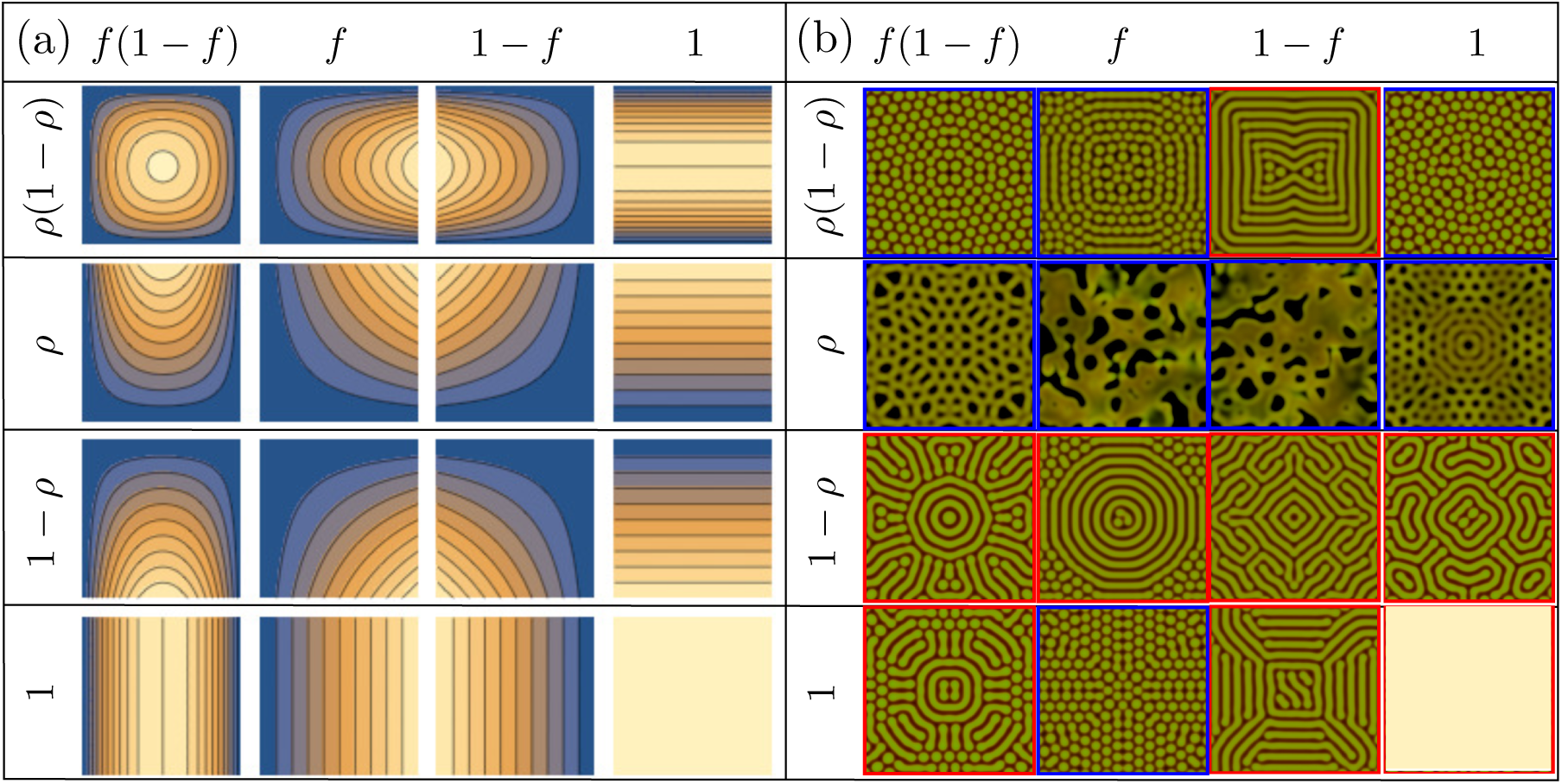
Patterns with various functional forms for defector’s diffusion coefficient. The density dependent functional form is determined by multiplying the functions in row and column. In (a), the shape of functions are shown in *f* (*x*-axis) and *ρ*(*y*-axis) space (contour plot). Blue and yellow colors represent low and high values at a given *f* and *ρ*, respectively. In (b), we present the patterns at a given functional forms for *r* = 2.32 and *σ* = 20. Without *ρ*, we observe striped patterns. Also, with 1 − *f* factor the striped patterns reemerge. The function *f* do exactly the opposite with 1 − *f* making dotted patterns. An uniform disk with densities *u* = *v* = 0.1 at a centre is used for an initial condition. Note that symmetry breaking for *r* = 2.28 and *D* = 4 comes from numerical underflow [26]. For the temporal evolution of all these patterns see Supplementary Video.

To avoid the threat of extinction, the cooperators fraction has to be enhanced. The behavior of (1 − *f*) is exactly boosting cooperators when cooperator’s fraction is too low. If defectors move fast when cooperator’s fraction *f* is small, *f* will increase. This results in population rescue and takes the system far from extinction, showing striped patterns.

We investigate the effect of diffusion driven by eco-evolutionary dynamics on pattern formation using numerical calculations. Mainly we have examined functions mimicking bacterial growth dynamics and human migration. These two functions show different patterns which lie at the ends of the survival spectrum: dotted and striped patterns. One behavior draws the system to a the edge of survival, whereas the other increases the size of the surviving regions. Furthermore, we estimate exactly what property of the diffusion dynamics makes it threaten extinction or conducive to survival. The results show that slow movement in low total population density threatens the system with extinction. However, population rescue is possible from the 1 − *f* behavior which is fast (slow) movement when the fraction of cooperators is low (high).

The equations of motion employed in this study are in principle modifications of the classical inhibitor-activator systems [42]. While it is clear that pattern formation is possible due to the higher diffusion coefficient of the inhibitor, we have provided a biologically meaningful reason for this diffusion disparity between activators and inhibitors. Across scales of organization it might be possible that defectors, cheaters, cancerous cells etc. have secured higher mobility as a benefit from not paying the costs of cooperation [39, 43].

Diffusion with a preference for forming alliances, can induce ecological conditions which are favourable to the spread of cooperation [44]. We have shown that diffusion driven by eco-evolutionary dynamics is instrumental in generating patterns which can be routinely seen in nature. Patterns in nature are generally expected to promote efficiency in organisms. Different organisms, with their different eco-evolutionary diffusion dynamics show different patterns which affect their ability to survive harsh conditions. Our finding may support the reason why we frequently observe dotted patterns in nature when the going gets tough [45]. On a macro scale this further encourages the use of both ecological as well as evolutionary approaches in understanding regular patterns observed widely in nature.

## Acknowledgements

We thank Christoph Hauert for comments and suggestions in improving the manuscript. We also thank David Rogers for improving the readability and clarity of the manuscript. Both authors acknowledge generous support from the Max Planck Society.

## Code availability

Appropriate computer code will be available upon request from the corresponding author.

### Appendix Average payoffs

In a public goods game, cooperators invest a fixed amount *c* in the common pool. The investments of all cooperators are amplified by multiplication factor *r* > 1, and then evenly returned to all individuals as the benefit. Under this setting, the payoffs for defectors and cooperators are given by,

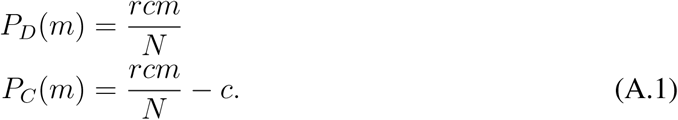

While we assume that the group size of the game is *N*, under eco-evolutionary dynamics it might be possible that there are not enough individuals to make up the group [46]. Therefore the actual group size can range from *S* = 2 to *N* while the rest is empty space. Individuals have a chance to meet and interact each other with a probability that is proportional to the total density in a well-mixed population. Therefore, the probability *p*(*S*; *N*) that an individual finds itself in a group of size *S* is given by,

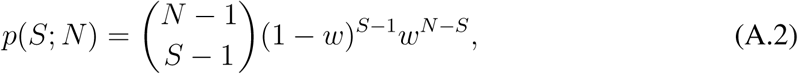

with sparseness *w* = 1 − *u* − *v*. The average payoffs for defectors and cooperators, *f_D_* and *f_C_*, are then calculated as follows,

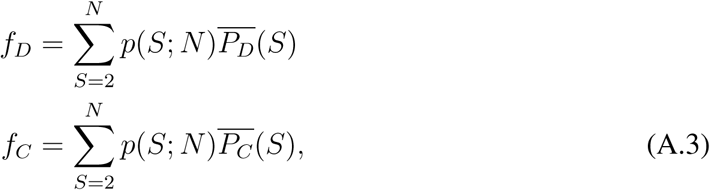

where
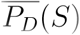
 and 
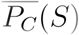
 are the expected payoffs for defectors and cooperators, respectively, with interacting group size *S* with the minimum group size being 2.

The probability that there are *m* cooperators among the *S* − 1 other individuals is given by *p_c_*(*m; S*),

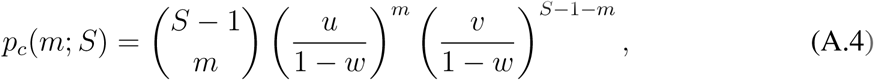

and the expected payoffs are written as,

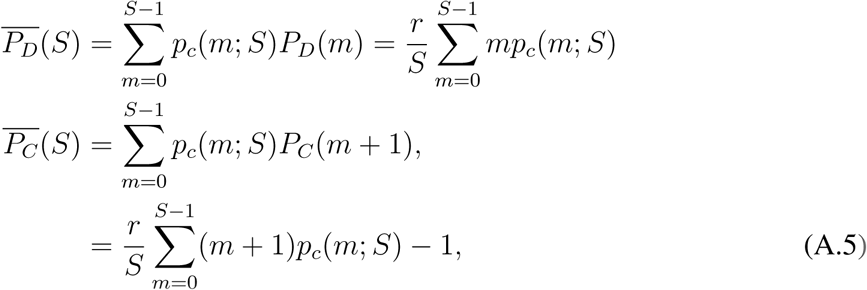

where we set the investment cost *c* = 1. Accordingly, the average payoffs *f_D_* and *f_C_* are calculated as follows,

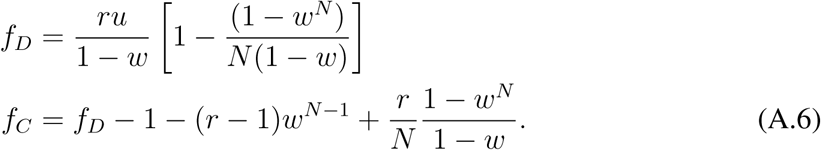

### Stable fixed point

Without spatial dynamics, the densities of cooperators and defectors change over time as described by the following equations:

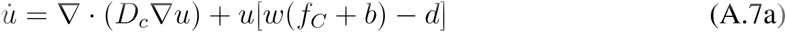

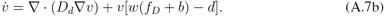

One of the fixed point(s) of this system is stable. Accordingly, the relation between visible *f* and *ρ* is imposed. Without spatial dynamics, we can perform a stability analysis for the system when we get an interior stable fixed point for *r* > *r_hopf_*. From the solution, we can get the relation between *f* and *ρ* at the stable fixed point. As shown in Fig. A.1, *f* and *ρ* are in a certain relationship i.e. the possible *f* and *ρ* pairs are restricted. When *ρ* increases *f* decreases along multiplication factor *r*.

Now we include spatial dynamics. With density dependent diffusion, the average *f* and *ρ* are in well agreement with the relationship established under no spatial effects for *r* > *r_hopf_*.

**Figure A.1:**
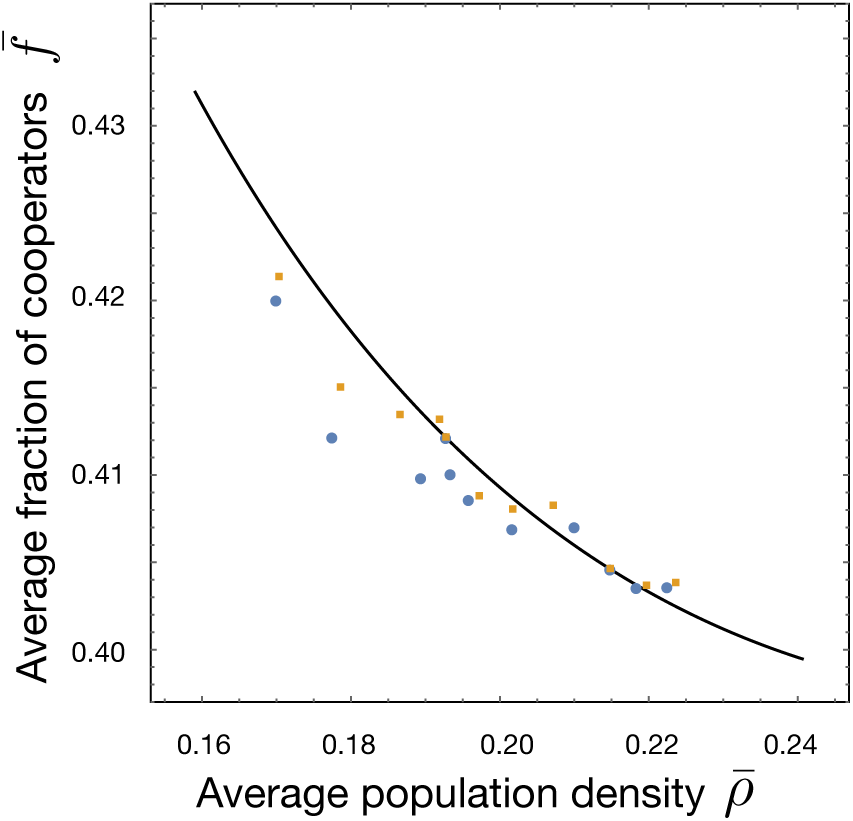
Stable fixed points of Eq. [A.7] for *r* > *r_hopf_*. The cooperator fraction *f* and total density *ρ* are in anti-correlation. When one increases, the other decreases. With density dependent diffusion, the average cooperator fraction 
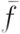
 and total density 
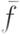
 keeps the relation. The points are the scatter plot of 
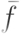
 and 
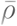
 with various *r*. Circles and squares indicate results with bacterial growth and human migration cases, respectively. The solid line is the fixed point evaluated numerically for *N* = 8, *d* = 1.2, *b* = 1 and *Ω* = 1 and *r* varying from 2:4 to 2:7 in steps of 0:001.

### Pattern analysis

Given the predominance of the two types of patterns, dotted and striped, we analyze the different properties between them for *r* < *r_hopf_*. Herein, we used configurations for *σ* = 18:75 and *r* = 2:36 with two different density dependent diffusion functions for analyzing patterns. Since *u* and *v* are inhomogeneous in *x, y* space, *D_d_*(*u, v*) is also spatially inhomogeneous.

Counterintuitively, the striped pattern has higher average cooperator fraction 
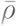
 than the dotted pattern (0:425 > 0:422). The average total density 
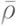
 of the striped pattern is smaller than that of the dotted pattern (0:156 < 0:159). It seems that the striped pattern is closer to the extinction phase because the high cooperator density and small total density, are the properties of patterns for small *r*. To understand this result, we examine the spatial distribution of *f* and *ρ*, because the average of all sub-populations may not be representative.

The two spatial dimensions *x, y* are both of size 283. We pick five *y* values to look at the spatial distribution of *u* and *v*, *y* = 10, 71, 141, 211, and 272. These are the values corresponding to regions near the domain boundaries, centre, and intermediate position (see Fig. A.2) respectively. We observe that the cooperator density is locally much higher than the defector density at the centre for the dotted pattern. The aforementioned does not appear in striped patterns. Hence, *f* is larger in the dotted pattern than the striped pattern at the centre. However, there are some places which have smaller *f* in the dotted pattern than the striped pattern, and thus 
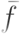
 in the dotted pattern is lower than that of the striped pattern even though they have a higher cooperator fraction locally.

**Figure A.2:**
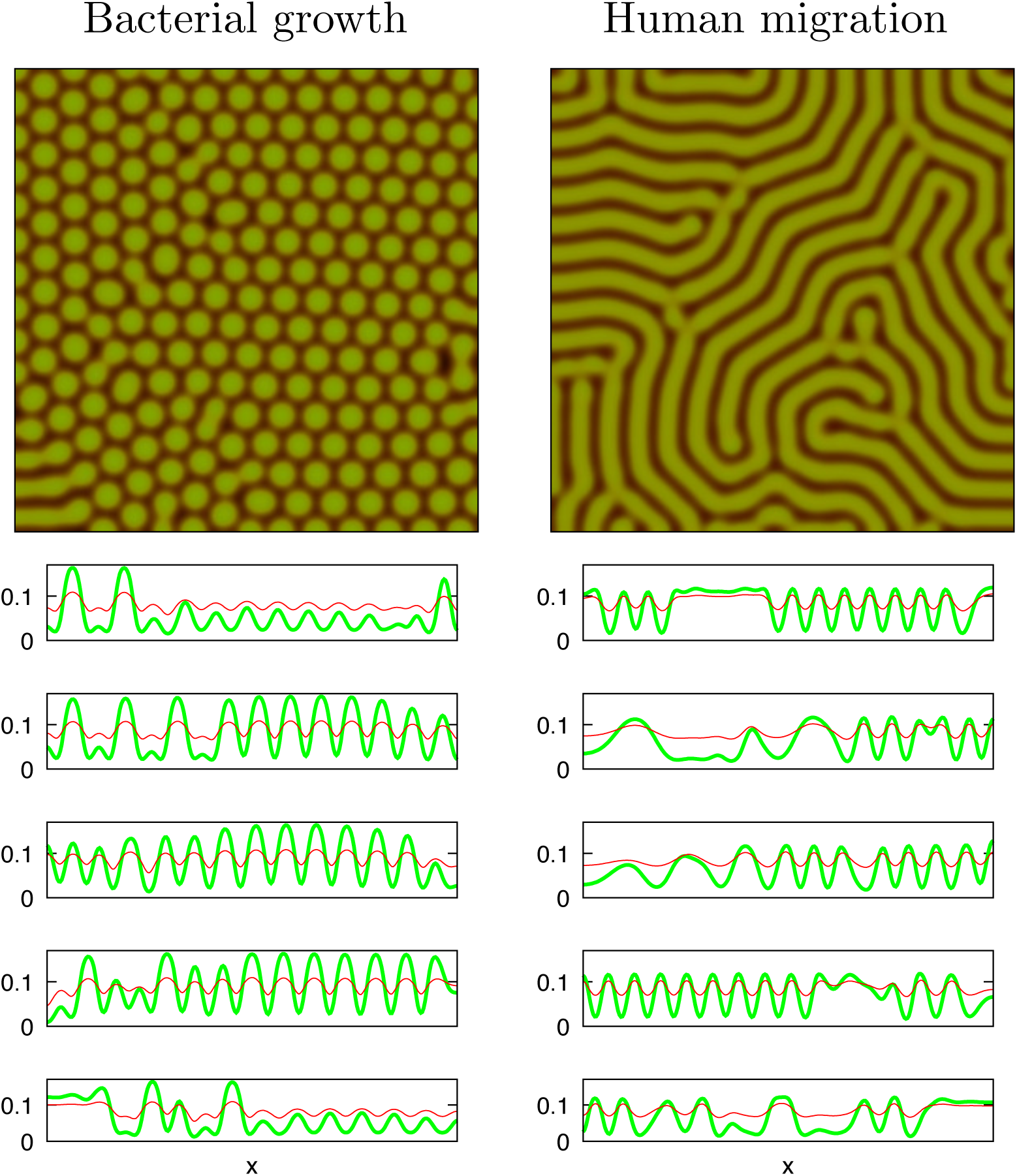
At a given pattern in each column, we horizontally slice along the *x*-axis at different points such as *y* = 271, 211, 141, 71, and 10 (from top to bottom) to show defector and cooperator denstities. This gives us the the densities *u* and *v* corresponding to regions near the domain boundaries (272,10), centre (141), and intermediate position (211,71). The left (right) column is for bacterial growth (human migration) case. Cooperator and defector densities are represented by thick green and thin red color, respectively

To see this effect, we look at the average cooperator fraction *f_y_* at each slice (average over *x* at a fixed *y*). As we can see in Fig. A.3, even though 
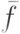
 for the dotted pattern is smaller than that of the striped pattern, locally *f* has a higher value. The dotted pattern has a much higher deviation of *f* and thus locally has much higher *f* than that of the striped pattern even though it has smaller average cooperator fraction 
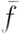
 It seems that the large fluctuations of *f* and *ρ* are the properties of the patterns close to the extinction phase.

**Figure A.3:**
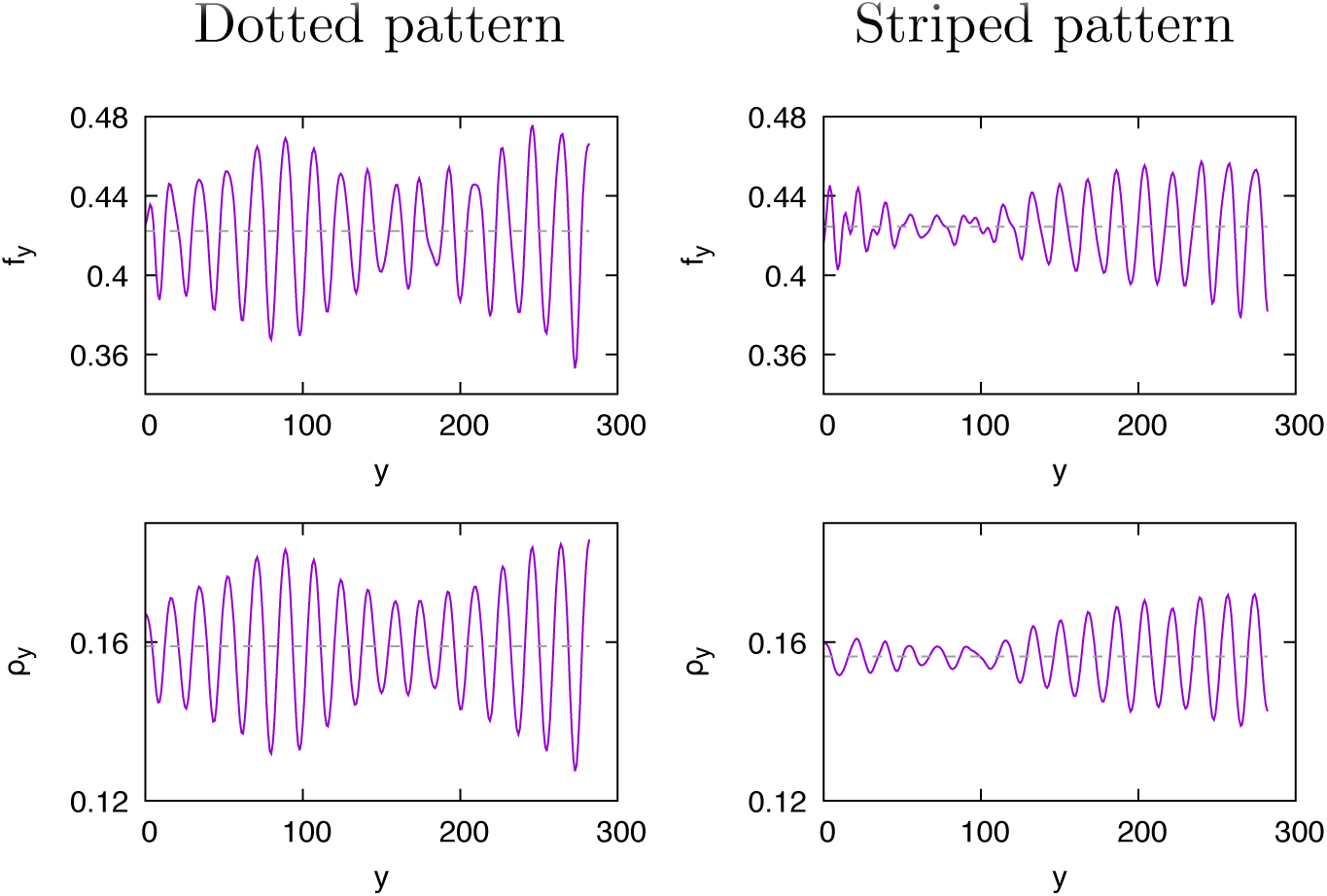
The average cooperator fraction and total density at *y*. The left column is the dotted pattern, while right one is the striped pattern. We see that *f_y_* and *ρ_y_*, are fluctuating in *y*, however the dotted pattern has a much larger deviation than the striped pattern Therefore, it locally has higher *f_y_* value.

### Color coding

For visualising cooperator and defector densities, we use green and red color for each type. The ratio of cooperator and defector determines the color spectrum. If only cooperators (defectors) are observed, the corresponding site is colored green (red). The yellow color appears when cooperators and defectors have the same density. The total density of the population is represented as the brightness of the color. For convenience, we use HSB color space which is a cylindrical coordinate (*r, θ, h*) = (saturation, hue, brightness). The angular variable *θ* represent color spectrum, hue where the red, green, and blue are located at *θ* = 0, *θ* = 2π/3 and *θ* = 4π/3, respectively. There is a relation between RGB and HSB coordinate, tan 
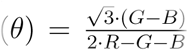
where *R, G*, and *B* are red, green, and blue coordinates of RGB space (cartesian coordinate).

The radius of circle *r* and height *h* indicate saturation and brightness, respectively. Since we do not use blue color, saturation becomes unity. Accordingly, brightness is used for the total population density *ρ* = *u* + *v*. If we use a linear function of *ρ* for brightness, it is hard to figure out the patterns for the small population density. For better visualisation, we formulate the brightness as

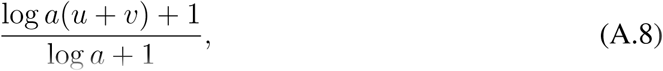

where a control parameter *a* (> - 1 and ≠ 0) determines the curvature of the brightness function in the total population density *ρ*(see Fig A.4).

**Figure A.4:**
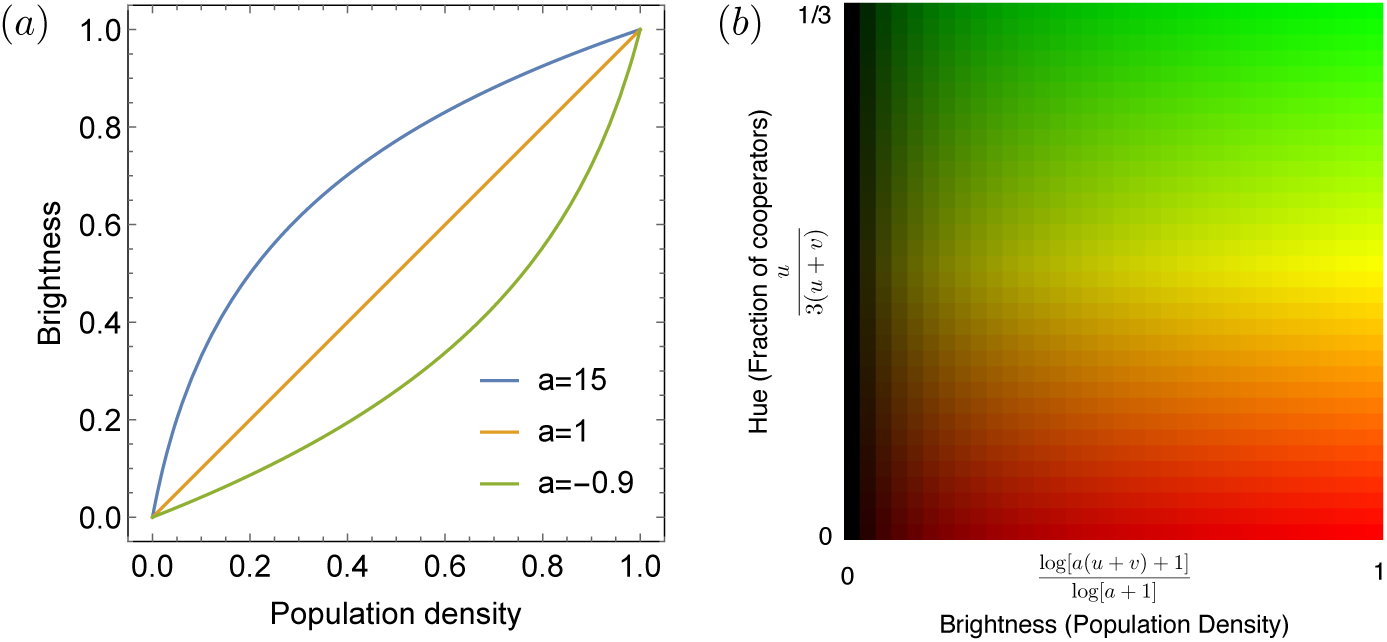
(a) To amplify brightness for small population density, we use non-linear relation between population density and brightness. When *a* = 1, the linear relation is recovered. For all figures, we use *a* = 15. (b) color for visualisation two densities (cooperator and defector). To show two components at the same site, we use color spectrum (hue) and brightness. Each represents a fraction of cooperators and total population density, respectively. Here, we normalise hue and brightness that each maximum becomes unity and use *a* = 15.

